## Supplementary material for "Local vulnerability and global connectivity jointly shape neurodegenerative disease propagation": S1 Table

| **Notation** | **Name** | **Epxression or value** | **Explanation** |
| --- | --- | --- | --- |
| $\Delta t$ | time step | $\Delta t=0.01$ | time increment in the simulations |
| $i$ | region label | $1\leq i\leq42$ | indexing regions |
| $(i, j)$ | edge label | $1\leq i, j\leq42$ | indexing edges |
| $N_{i}$ | normal population  in region $i$ | N/A | total number of normal agents in region $i$ |
| $M_{i}$ | misfolded population  in region $i$ | N/A | total number of misfolded agents in region $i$ |
| $N_{(i, j)}$ | normal population  in edge $(i,j)$ | N/A | total number of normal agents in edge $(i, j)$ |
| $M_{(i,j)}$ | misfolded population  in edge $(i,j)$ | N/A | total number of misfolded agents in edge $(i, j)$ |
| $\alpha_{i}$ | synthesis rate in region $i$ | $\alpha_{i}=\Phi_{0,1}(SNCA_{i})$ where $SNCA_{i}$ is the SNCA expression (z-score) in region $i$ | the probability that a new normal agent gets synthesized in each voxel of region $i$ per unit time |
| $\beta_{i}$ | clearance rate in region $i$ | $\beta_{i}=\Phi_{0,1}(GBA_{i})$ where $GBA_{i}$ is the GBA expression (z-score) in region $i$ | the probability that an existing agent (either normal or misfolded) in region $i$ gets cleared per unit time |
| $S_{i}$ | region size | N/A | voxel counts of region $i$ |
| $\gamma_{i}^{0}$ | baseline transmission rate | $\gamma_{i}^{0}=1/S_{i}$ | the probability for a single misfolded agent to transmit the disease to other agents per unit time |
| $\gamma_{i}$ | transmission probability | $1-\exp(M_{i}\ln(1-\gamma_{i}^{0}))$ | the probability that normal agents get infected (by at least one of the misfolded agents) per unit time |
| $w_{ij}$ | connection strength of edge $(i, j)$ | normalized fiber tracts density between region $i$ and $j$ | determining the probability of entering edge $(i, j)$ when exiting region $i$ per unit time |
| $l_{ij}$ | edge length of edge $(i, j)$ | mean length of fiber tracts between region $i$ and $j$ | determining the probability of exiting edge $(i, j)$ per unit time |
| $\rho_{i}$ | the probability of remaining in region $i$ | $\rho_{i}=0.5$ for all $i$ | agents in region $i$ have equal probability of remaining in region $i$ or exiting region $i$ per unit time |
| $\text{f}\text{c}_{ij}$ | functional connectivity of edge $(i,j)$ | N/A | biasing agents toward regions showing greater co-activation pattern |
| $k$ | weight of functional connectivity | N/A | controlling the influence of functional connectivity in driving disease spread |
| $k_{1}$ | weight of atrophy accrual due to accumulation of misfolded agents | $k_{1}+k_{2}=1$ | controlling the contribution of native misfolded agents to total atrophy growth |
| $k_{2}$ | weight of atrophy accrual due to deafferentation | $k_{1}+k_{2}=1$ | controlling the contribution of deafferentation to total atrophy growth |
| $r_{i}(t)$ | the ratio of misfolded agents in region $i$ | $r_{i}\left( t \right)=\frac{M_{i}\left( t \right)}{N_{i}\left( t \right)+M_{i}\left( t \right)}$ | measuring the burden of misfolded agents in region $i$ at time $t$ |
