## Supplementary material for "Local vulnerability and global connectivity jointly shape neurodegenerative disease propagation": S1 Text

Analysis of the fixed points

Although there is no analytical solution for α-synuclein concentration, phase plane analysis is helpful in finding fixed points of the system. Considering that the rates of incoming and outgoing agents in the edges (axons) are equal when the system is at the stable point, and that no clearance, synthesis, or misfolding occurs in the edges, the effects of the propagation process are negligible in the analysis of the system's fixed points. Therefore, we sought to use the “overall” or “total’’ synthesis rate ($\alpha$), normal agent clearance rate ($\beta_{1}$), misfolded agent clearance rate ($\beta_{2}$), and transmission rate ($\gamma$) to approximately analyze the entire system using a series of differential equations (equations (S1)-(S3) below). Likewise, we use $N$, $M$ to represent the total population of normal and misfolded agents in the entire brain. $\beta_{1}$, $\beta_{2}$, $\gamma$ depend on the actual $N_{i}$, $M_{i}$ and thus are not static values (to see this, for example, the total cleared normal agents per unit time is $\sum_{i} \beta_{i}N_{i}$, so the “overall” clearance rate $\sum_{i} \beta_{i}N_{i}/N$ depends on real-time $N_{i}$; it is not the “real” clearance rate but an approximation of the total rate of clearance); the total synthesis rate $\alpha=\sum_{i} \alpha_{i}S_{i}$, where $\alpha_{i}$ is the empirical synthesis rate in region $i$ in the formal model and $S_{i}$ is region size. Note that the actual spreading dynamics cannot be fully described by the following differential equations (S1)-(S3). However, they are helpful in analyzing the possible states of disease propagation.

**Growth of Normal α-synuclein.** The system of normal α-synuclein growth approximates like:

$$\begin{aligned} \frac{\text{d}N}{\text{d}t}=\alpha- \beta_{1}N \#\#(S1) \end{aligned}$$

with $\alpha\gg1$, ${0<\beta}_{1}<1$, obviously $N=\alpha/\beta_{1}$ is a stable point (which is the baseline density of endogenous α-synuclein, Fig 3, blue curve).

**Spread of misfolded α-syn.** The system with misfolded agents injected behaves like:

$$\begin{aligned} \frac{\text{d}N}{\text{d}t}=\alpha-\beta_{1}N-\left( 1-\beta_{1} \right)\left( 1-\left( 1-\gamma\right)^{M} \right)N\#\left( S2 \right) \end{aligned}$$

$$\begin{aligned} \frac{\text{d}M}{\text{d}t}=\left( 1-\beta_{1} \right)\left( 1-\left( 1-\gamma\right)^{M} \right)N- \beta_{2}M\#\left( S3 \right) \end{aligned}$$

The nullclines of $N$, $M$ are

$$\begin{aligned} M=\frac{\ln\left( \frac{N-\alpha}{N\left( 1-\beta_{1} \right)} \right)}{\ln\left( 1-\gamma\right)}\#\left( S4 \right) \end{aligned}$$

$$\begin{aligned} N=\frac{\beta_{2}M}{\left( 1-\beta_{1} \right)\left( 1-\left( 1-\gamma\right)^{M} \right)}\#\left( S5 \right) \end{aligned}$$

Note equation (S5) is not defined at $M=0$. To study the number of fixed points and their positions, we need to determine the number of intersections of the two nullclines (S4) and (S5), and where they intersect. Adding (S2) and (S3), it is easy to see $\left( N, M \right)=(\alpha/\beta_{1},0)$ is one fixed point. It is easy to find that $M$ decreases monotonously with $N$ in (S4) and passes $(\alpha/\beta_{1},0)$. Therefore, the monotony and position of line (S5) relative to line (S4) becomes crucial. To find the monotony of (S5), we take its first order derivative

$$\begin{aligned} N^{'}=\frac{\beta_{2}\left( 1-\beta_{1} \right)\left( \left( 1-\left( 1-\gamma\right)^{M} \right)+M\left( 1-\gamma\right)^{M}\ln\left( 1-\gamma\right) \right)}{\left( \left( 1-\beta_{1} \right)\left( 1-\left( 1-\gamma\right)^{M} \right) \right)^{2}}\#\left( S6 \right) \end{aligned}$$

When $M=0$, the first order derivative is 0. We then take the second derivative

$$\begin{aligned} N^{''}= C\left( \ln\left( 1-\gamma\right) \right)^{2}\left( 1-\gamma\right)^{M}M\#\left( S7 \right) \end{aligned}$$

where $C$ is a positive constant. When $M>0$, (S7) is positive thus the first order derivative (S5) increases monotonously with $M$, and when $M<0$, (S5) decreases monotonously with $M$. Therefore, (S6) is positive hence in (S5), $N$ increases monotonously with $M$, namely, there can be up to one intersection (denoted by ${(N}^{*}, M^{*})$ in the following) of the two nullclines apart from $\left( N, M \right)=(\alpha/\beta_{1},0)$. It determines that the system can have up to two fixed points, one is $(\alpha/\beta_{1},0)$, and the other ${(N}^{*}, M^{*})$, which has no closed-form expression.

It is also important to determine the position of ${(N}^{*}, M^{*})$, because $M^{*}<0$ is not realistic in the actual disease spread. Under this condition, an outbreak can never take place. Taking the limit of (S5) $\lim_{M\to0} N=\beta_{2}/\left( \beta_{1}-1 \right)\ln(1-\gamma)$, we can see the intercept of (S5) on the $N$ axis is $\beta_{2}/(\beta_{1}-1)ln(1-\gamma)$. When (i) $\beta_{2}=\left( \beta_{1}-1 \right)\ln\left( 1-\gamma\right)\alpha/\beta_{1}$, $N^{*}=\alpha/\beta_{1}$, i.e., ${(N}^{*}, M^{*})$ “merges” with $(\alpha/\beta_{1},0)$; (ii) when $\beta_{2}>\left( \beta_{1}-1 \right)\ln\left( 1-\gamma\right)\alpha/\beta_{1}$, $N^{*}>\alpha/\beta_{1}$, $M^{*}<0$ which is not realistic; (iii) when $\beta_{2}<\left( \beta_{1}-1 \right)\ln\left( 1-\gamma\right)\alpha/\beta_{1}$, $N^{*}<\alpha/\beta_{1}$, $M^{*}>0$. Our results (with full spread in the end) are based on (iii).

To study why different choices of seed region and injected α-synuclein may lead to different states (extinction or outbreak), we also investigated under what conditions the fixed points are stable. This can be studied by taking the Jacobian matrix of the system linearized around the fixed points:

$$\begin{aligned} J=\left( \begin{matrix} -\beta_{1}-\left( 1-\beta_{1} \right)\left( 1-\left( 1-\gamma\right)^{M} \right) & \left( 1-\beta_{1} \right)\left( 1-\gamma\right)^{M}\ln\left( 1-\gamma\right)N \\ \left( 1-\beta_{1} \right)\left( 1-\left( 1-\gamma\right)^{M} \right) & -\beta_{2}-\left( 1-\beta_{1} \right)\left( 1-\gamma\right)^{M}\ln\left( 1-\gamma\right)N \end{matrix} \right)\#\left( S8 \right) \end{aligned}$$

At $(\alpha/\beta_{1},0)$, $J\left. \right|_{(\alpha/\beta_{1}, 0)}=\left( \begin{matrix} -\beta_{1} & \left( 1-\beta_{1} \right)\ln\left( 1-\gamma\right)\alpha/\beta_{1} \\ 0 & -\beta_{2}-(1-\beta_{1})ln(1-\gamma)\alpha/\beta_{1} \end{matrix} \right)$ has eigenvalues $-\beta_{1}$ and $-\beta_{2}-(1-\beta_{1})ln(1-\gamma)\alpha/\beta_{1}$.

When $\beta_{2}>\left( \beta_{1}-1 \right)\ln\left( 1-\gamma\right)\alpha/\beta_{1}$, both eigenvalues are negative hence $(\alpha/\beta_{1},0)$ is stable (disease extinction).

The injection of misfolded agents introduces a small perturbation to the system at $(\alpha/\beta_{1},0)$. As the choice of seed region and injection amount affect the approximations of $\beta_{1}$, $\beta_{2}$, $\gamma$ in equation (S2) and (S3) that are used to analyze the system, the disease will either die out or fully spread. It is more difficult to initiate the disease spread in seed regions with relatively large $\beta_{1}$, $\beta_{2}$ and small $\gamma$ (i.e., more resistant to disease spread), as it is more likely to satisfy the condition $\beta_{2}>\left( \beta_{1}-1 \right)\ln\left( 1-\gamma\right)\alpha/\beta_{1}$ initially. As mentioned before, $\beta_{1}$, $\beta_{2}$, $\gamma$ are not static and depend on the real-time $N_{i}$, $M_{i}$s; at the other fixed point ${(N}^{*}, M^{*})$, the parameter set is not the same as the one near $(\alpha/\beta_{1},0)$. Therefore it is also possible, in theory, that certain choices of parameters may lead to an outbreak followed by gradual extinction. S1 Fig gives an example of the two fixed points under $\alpha=5000$, $\beta_{1}=0.5\{$1}, $\beta_{2}=0.5$, $\gamma=0.001$ in which the nullclines of $N$ and $M$ intersects at $(5017.15, 4982.85)$ and $(10000, 0)$.
